# Dynamic changes in neuronal and glial GAL4 driver expression during *Drosophila* ageing

**DOI:** 10.1101/2024.06.16.599238

**Authors:** Caroline Delandre, John P. D. McMullen, Owen J. Marshall

## Abstract

Understanding how diverse cell types come together to form a functioning brain relies on the ability to specifically target these cells. This is often done using genetic tools such as the GAL4/UAS system in *Drosophila melanogaster*. Surprisingly, despite its extensive usage during studies of the ageing brain, detailed spatio-temporal characterisation of GAL4 driver lines in adult flies has been lacking. Here we show that three commonly used neuronal drivers (*elav[C155]-GAL4*, *nSyb[R57C10]-GAL4* and *ChAT-GAL4*) and the commonly used glial driver *repo-GAL4* all show rapid and pronounced decreases in activity over the first 1.5 weeks of adult life, with activity becoming undetectable in some regions after 30 days. In addition to an overall decrease in GAL4 activity over time, we found notable differences in spatial patterns, mostly occurring soon after eclosion. Although all lines showed these changes, the *nSyb-GAL4* line exhibited the most consistent and stable expression patterns over ageing. Our findings suggest that gene transcription of key loci decreases in the aged brain, a finding broadly similar to previous work in mammalian brains. Our results also raise questions over past work on long-term expression of disease models in the brain, and stress the need to find better genetic tools for ageing studies.

## Introduction

Ever since the intricate drawings by Santiago Ramón y Cajal over a century ago, the wide diversity of cell types populating the nervous system has presented a formidable challenge to neuroscientists (Ramón y Cajal, 1904). Understanding how this diversity is formed and how the different cell types within the brain change in response to ageing and disease is a goal that the genetic tools of *Drosophila melanogaster* have made more attainable. The prime example of this is the GAL4/UAS bipartite gene expression system, in which the cell type-specific expression of the yeast transcription factor GAL4 can drive expression of a transgene (Brand and Perrimon, 1993). Thousands of GAL4 drivers are now available to the fly community, enabling the targeting of transgene expression to cell types and subtypes, sometimes at the resolution of a single neuron within the whole brain (Jenett et al., 2012). As most fly research has become heavily reliant on GAL4 lines, it is critically important to carefully describe their expression pattern, both spatial and temporal. Indeed, the chromatin environment of enhancers driving GAL4 expression can often be highly dynamic throughout development, and therefore, a specific GAL4 driver may target different cell types over time (Markstein et al., 2008). This characterisation is particularly important in the field of ageing, as there is a knowledge gap in our understanding of the activity behaviour of commonly used nervous system GAL4 lines in older flies.

Anecdotal reports in the literature have indicated potential problems with widely used GAL4 lines. The most prominent is the developmental expression in neuroblasts and glia of the *elav[C155]-GAL4* line, long considered to be restricted to post-mitotic neurons and a popular choice for pan-neuronal expression (Berger et al., 2007). Other groups also found this same driver and other neuronal ones to be expressed outside of the nervous system during part of development or in adult flies (Casas-Tintó et al., 2017; Weaver et al., 2020; Winant et al., 2024). However, most reports examine expression patterns only in development or young adults, and we were surprised by the small number of reports describing how GAL4 drivers change as the brain ages.

That changes in driver expression can occur during ageing is clear. Early work characterising enhancer traps using beta-galactosidase already showed changes in expression patterns across lifespan (Helfand et al., 1995). These were followed by several reports from the Benzer and Seroude groups who found expression changes of a GAL4 enhancer trap set across lifespan in whole flies (Poirier et al., 2008; Seroude, 2002; Seroude et al., 2002).

Another caveat is that a common readout for GAL4 driver expression is a stable fluorescent reporter driven through development. When the tissue is imaged, fluorescence levels result from the protein accumulation over the lifetime of the organism, thereby masking the actual expression dynamic of the GAL4 driver. This is also often followed by immunostaining against induced reporter proteins, which can reduce the signal-to-noise ratio of fluorescent reporters and further mask region-specific expression differences (Wissing et al., 2022). Furthermore, the insertion site of reporter transgenes may also play a role in determining the final readout, and it is important to ensure its chromatin environment does not vary significantly throughout development or ageing (Pfeiffer et al., 2010).

Finally, several studies have used RNA sequencing to investigate gene expression levels in the ageing brain, both in mammals and flies. These studies have suggested that gene expression of some genes may decrease or increase in old brains (Ham and Lee, 2020; Pacifico et al., 2018). However, as RNA sequencing measures only relative abundance of a transcript rather than absolute abundance, the exact nature of gene expression changes during ageing remains unclear (Lovén et al., 2012). When measuring total RNA as flies age, older studies found a decline in RNA levels, especially steep within the first days after eclosion (Tahoe et al., 2004). This was postulated to be due to the transition between metamorphosis, requiring high gene expression levels, and adulthood (Shikama and Brack, 1996). More recently, a similar early decline was reported in the fly brain using single-cell transcriptomics (Davie et al., 2018; Lu et al., 2023).

Given all the above factors and the fact that *Drosophila* is becoming a powerful model organism to study the fundamental mechanisms of ageing and neurodegenerative diseases (for review, see McGurk et al., 2015), we sought to profile the expression dynamics and distribution of common brain GAL4 drivers in older flies.

Here, we used the TARGET system to temporally restrict GAL4 driver activity and profile the expression of commonly used neuronal and glial driver lines during adult brain ageing: *elav[C155]-GAL4* and *nSyb-GAL4* (also known as *R57C10-GAL4*), drivers widely used as pan-neuronal, *ChAT-GAL4*, a cholinergic neuronal driver, and the pan-glial driver *repo-GAL4*. We found that their expression patterns in young adult brains were surprisingly not as uniform as expected, instead showing enrichment in particular brain regions. More importantly, we found that there was a dramatic decrease in expression during the first days after eclosion, with expression continuing to decrease throughout ageing.

## Materials and method

### *Drosophila* maintenance and stocks

Flies were cultured in a standard food medium consisting of corn meal (35 g/L), dextrose (55 g/L), and yeast (50 g/L), and kept at 18°C or 25°C in incubators with controlled humidity (70%) and a light:dark cycle of 12 hours each. Drivers used were *P{GawB}elav[C155]* (*elav[C155]-GAL4*; Bloomington #458), *P{GMR57C10-GAL4}attP2* (*nSyb-GAL4*; Bloomington #39171), *Mi{Trojan-GAL4.0}ChAT[MI04508]* (*ChAT-GAL4*; Bloomington #60317), and *P{GAL4}repo* (*repo-GAL4*; Bloomington #7415). GFP stocks were *10XUAS-IVS-myr::GFP* in attP2 (Bloomington #32197), attP40 (Bloomington #32198), or su(Hw)attP5 (Bloomington #32199). Temperature-sensitive GAL4 inhibition was achieved using *P{tubP-GAL80[ts]}* either on chromosome 2 or 3 and keeping flies at 18°C. Flies were transferred to a 29°C incubator for 24 hours to induce expression of UAS reporters.

### Brain dissection and imaging

Adult brains were dissected in phosphate-buffered saline (PBS) and fixed in 4% paraformaldehyde, followed by three washes in PBS + 0.3% Triton X-100 and mounting onto slides with VECTASHIELD PLUS with DAPI (Vector Laboratories, USA). Slides were imaged with an FV3000 confocal laser scanning microscope (Olympus, Japan) under a 20X objective. Optimal laser settings were independently adjusted for each GAL4 driver to ensure proper spatial resolution. However, imaging settings were not changed when comparing time points. Four to eight brains were dissected for each time point and driver (except for *ChAT-GAL4* at 15 DPE with only three brains). Since *elav[C155]-GAL4* is located on the X chromosome, only brains from male flies were dissected.

### Analysis

Images were analysed using the Icy open source imaging software (de Chaumont et al., 2012) by creating regions of interest in selected brain areas and recording the GFP mean pixel intensity. Values were then plotted using R (R Core Team, 2021).

## Results

### Commonly used neuronal GAL4 drivers show differential spatial expression patterns in young adult brains

To test the reliability of brain GAL4 drivers throughout adult life, we started by investigating the expression profile of three commonly used neuronal drivers in young adult brains. The classic pan-neuronal *elav[C155]-GAL4* is an enhancer trap line where GAL4 was inserted in the *elav* promoter region (Ogienko et al., 2020). *nSyb-GAL4* was generated by cloning a fragment from the *nSyb* gene upstream of GAL4 and inserting the resulting plasmid into attP2 (chromosome 3) (Pfeiffer et al., 2008). *ChAT-GAL4* was engineered by inserting a GAL4 Trojan cassette into a MiMIC integration site located in an intron of the *ChAT* gene. This cassette places a T2A-GAL4 fragment flanked by splice acceptors so the GAL4 expression should reproduce the native pattern of *ChAT* (Diao et al., 2015). We obtained a baseline of their spatial expression profile by crossing these drivers to *UAS-myrGFP* (inserted in attP2) and culturing the progeny at 25°C until one day post-eclosion (DPE) when they were dissected. The GFP expression pattern of both pan-neuronal drivers was quite different from what we expected; instead of a uniform signal present across the brain, we observed a clear enrichment in specific areas, most notably the mushroom body (Fig. 1A). Between the two drivers, *elav[C155]-GAL4* showed the most dramatic differential expression, with very low signal in the optic lobes and midbrain areas other than the mushroom body (especially the alpha/beta lobes) and the antennal lobes. *ChAT-GAL4* also showed a mushroom body bias, but otherwise seemed to replicate the expression of cholinergic neurons (Salvaterra and Kitamoto, 2001).

**Figure 1.**
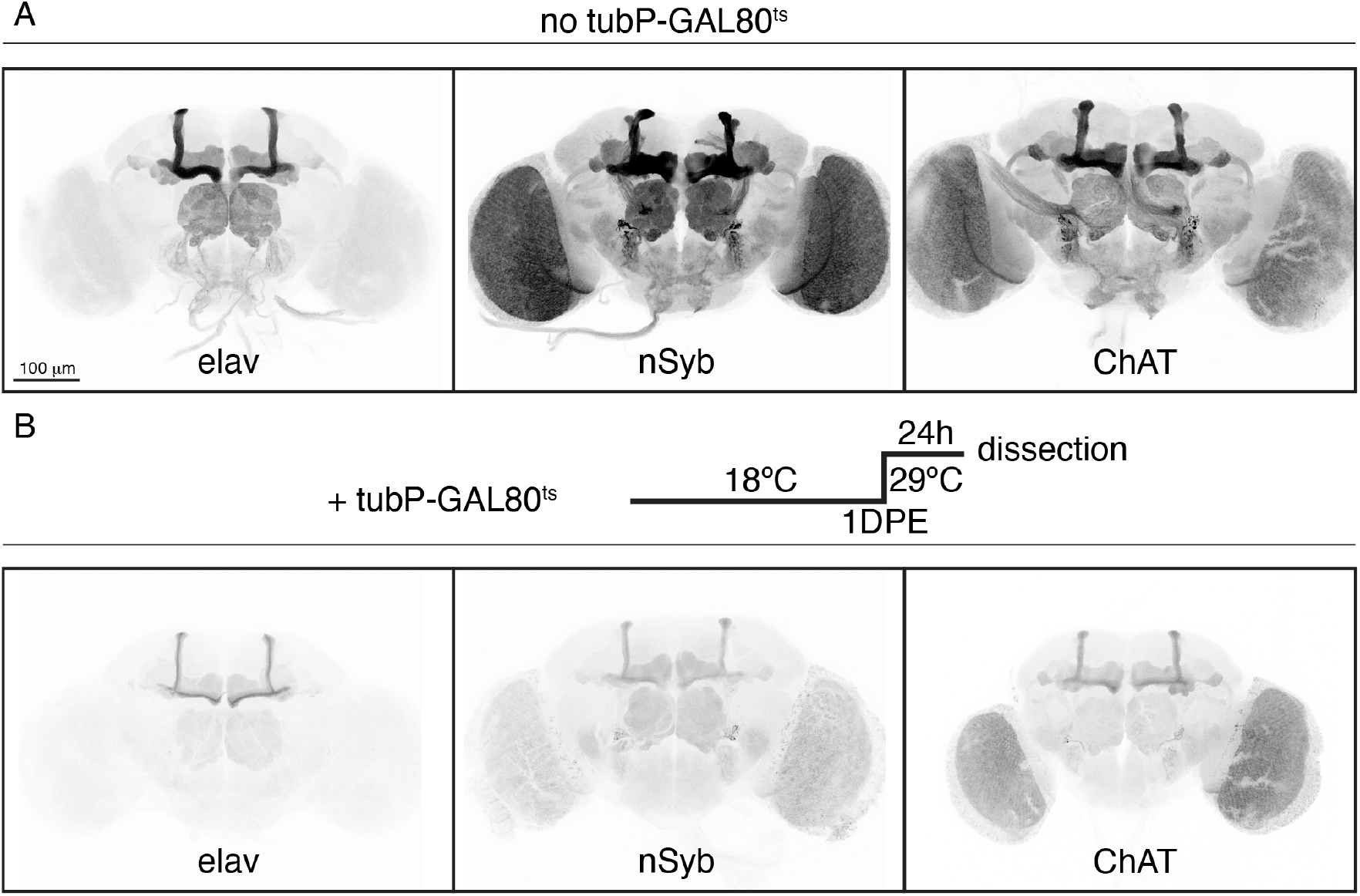
Common neuronal GAL4 drivers are expressed differentially across the brain. (A) 1-DPE brains were imaged for membrane-bound GFP driven by *elav[C155]-GAL4*, *nSyb[R57C10]-GAL4*, or *ChAT-GAL4*, either throughout development (A) or for 24 hours right before dissection via a temperature-sensitive GAL80 (B).

We then investigated how temporally restricting driver activity would affect their expression profiles by adding a temperature-sensitive GAL80 under the promoter of *alphaTub84B* (*tubP-GAL80[ts]*). Flies were reared at 18°C until 1 DPE when brains were placed at 29°C for 24 hours to induce GFP expression. As opposed to driving GFP expression throughout development, this technique should prevent background from accumulation of GAL4 and GFP molecules produced before 1 DPE. Indeed, when compared with the previous images, the brains that were induced at 1 DPE showed a decrease in overall GFP signal (Fig. 1B). The expression profile in these images thus represents the actual activity of these drivers at 1 DPE. Spatial expression profiles were overall similar to images from brains that were not temporally restricted, indicating these GAL4 lines are likely driving expression in the same subset of neurons during the few days before eclosion. Taken together, these results show that both pan-neuronal drivers do not drive expression ubiquitously across neuronal cell types and that temporally restricting their activity provides a more accurate representation of GAL4 activity levels at that time point.

### Neuronal driver activity decreases throughout adult life in a brain region-specific manner

We then asked whether the expression profiles of neuronal drivers change as brains age. Using *tubP-GAL80[ts]*, we restricted GAL4 activity to nine 24-hour time windows, from 1 to 90 DPE (Fig. 2). We chose four representative brain regions for quantification: antennal lobe, the α and γ lobes of the mushroom body, and optic lobe (Fig. 2B). All three neuronal drivers exhibited striking declines in activity. For *elav[C155]-GAL4*, we noticed a reduction in expression profile starting at just 5 DPE in the mushroom body alpha/beta lobes. The mushroom body alpha’/beta’ lobes together with the optic lobes became barely detectable from 5 to 10 DPE. Moreover, there was a gradual decrease in overall driver activity, notably during the first 30 days.

**Figure 2.**
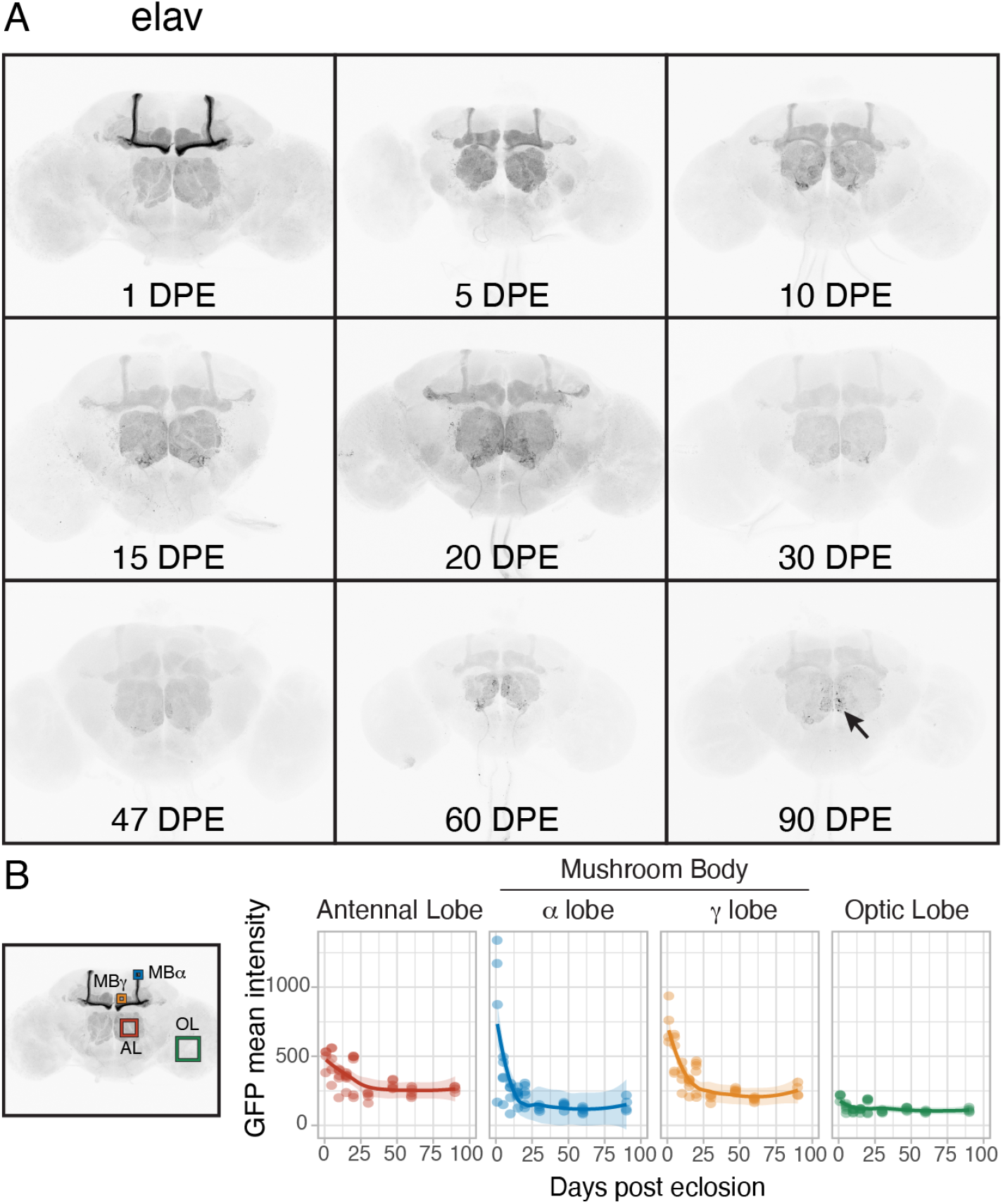
(A) Flies expressing membrane-bound GFP under the control of *elav[C155]-GAL4* and a temperature-sensitive GAL80 were induced for 24-hour time windows by shifting from 18°C to 29°C at nine time points throughout adult life (up to 90 DPE). Arrow indicates the appearance of individual neurons in older brains. (B) Four brain regions were chosen for quantification of GFP mean intensity (n = 4-8).

Similarly, there was a gradual decline, albeit not as steep, in *nSyb-GAL4* activity during the first 30 days of adulthood (Fig. 3). There was also a reduction in the signal difference between the alpha/beta and gamma lobes: from 30 DPE onwards, the beta lobes were almost indistinguishable from the gamma lobes. The expression in the optic lobes gradually lost its uniform pattern as brains aged, with GFP aggregates most obvious in the oldest brains. Interestingly, a small group of neurons located next to the antennal lobes remained strongly labelled throughout the experiment. *ChAT-GAL4*, but not *elav[C155]-GAL4*, also drove persistent expression in this neuronal subset (Fig. 4A), ruling out a potential artefact specific to the *nSyb-GAL4* construct design.

**Figure 3.**
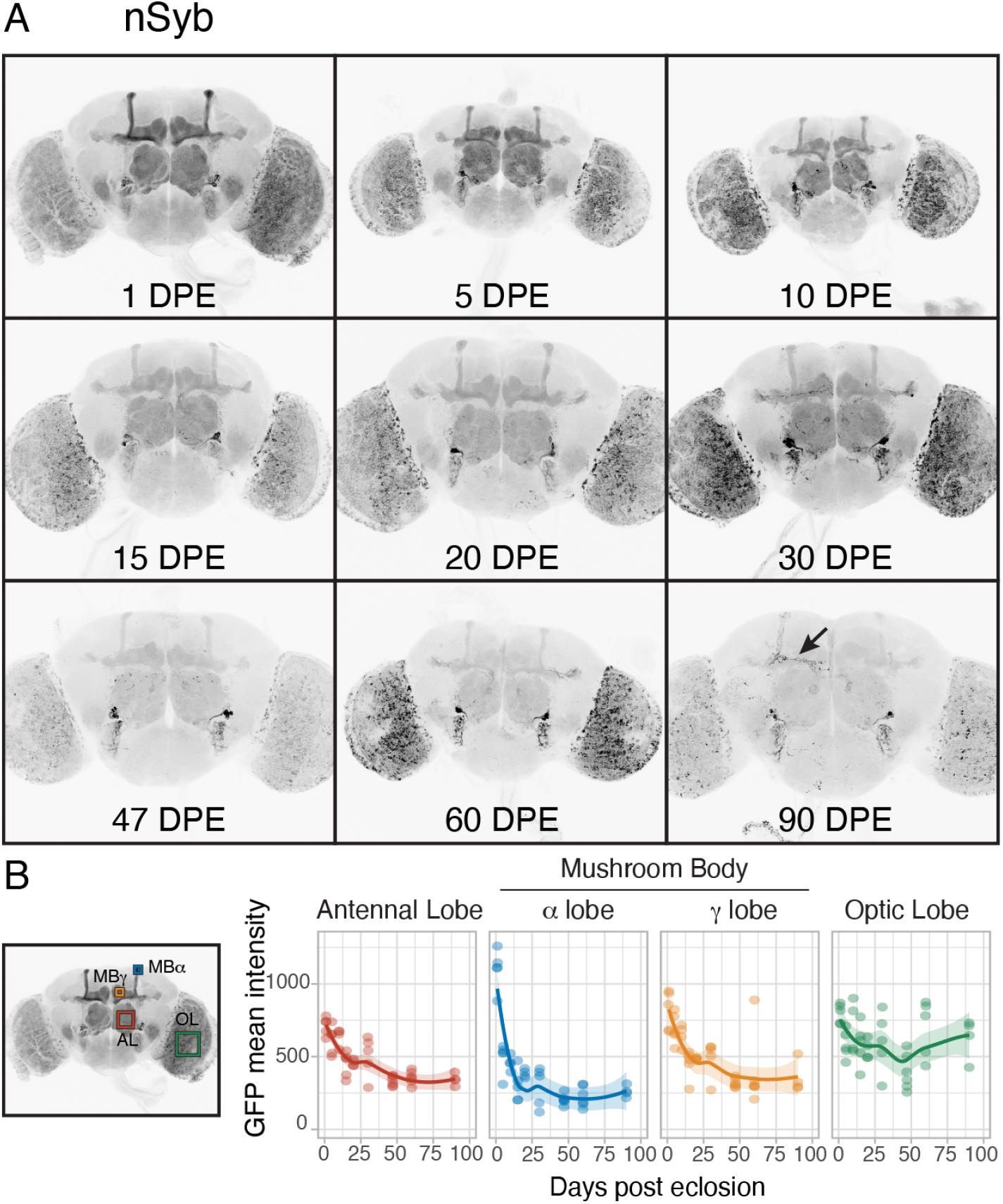
(A) Flies expressing membrane-bound GFP under the control of *nSyb[R57C10]-GAL4* and a temperature-sensitive GAL80 were induced for 24-hour time windows by shifting from 18°C to 29°C at nine time points throughout adult life (up to 90 DPE). Arrow indicates the appearance of individual neurons in older brains. (B) Four brain regions were chosen for quantification of GFP mean intensity (n = 4-8).

**Figure 4.**
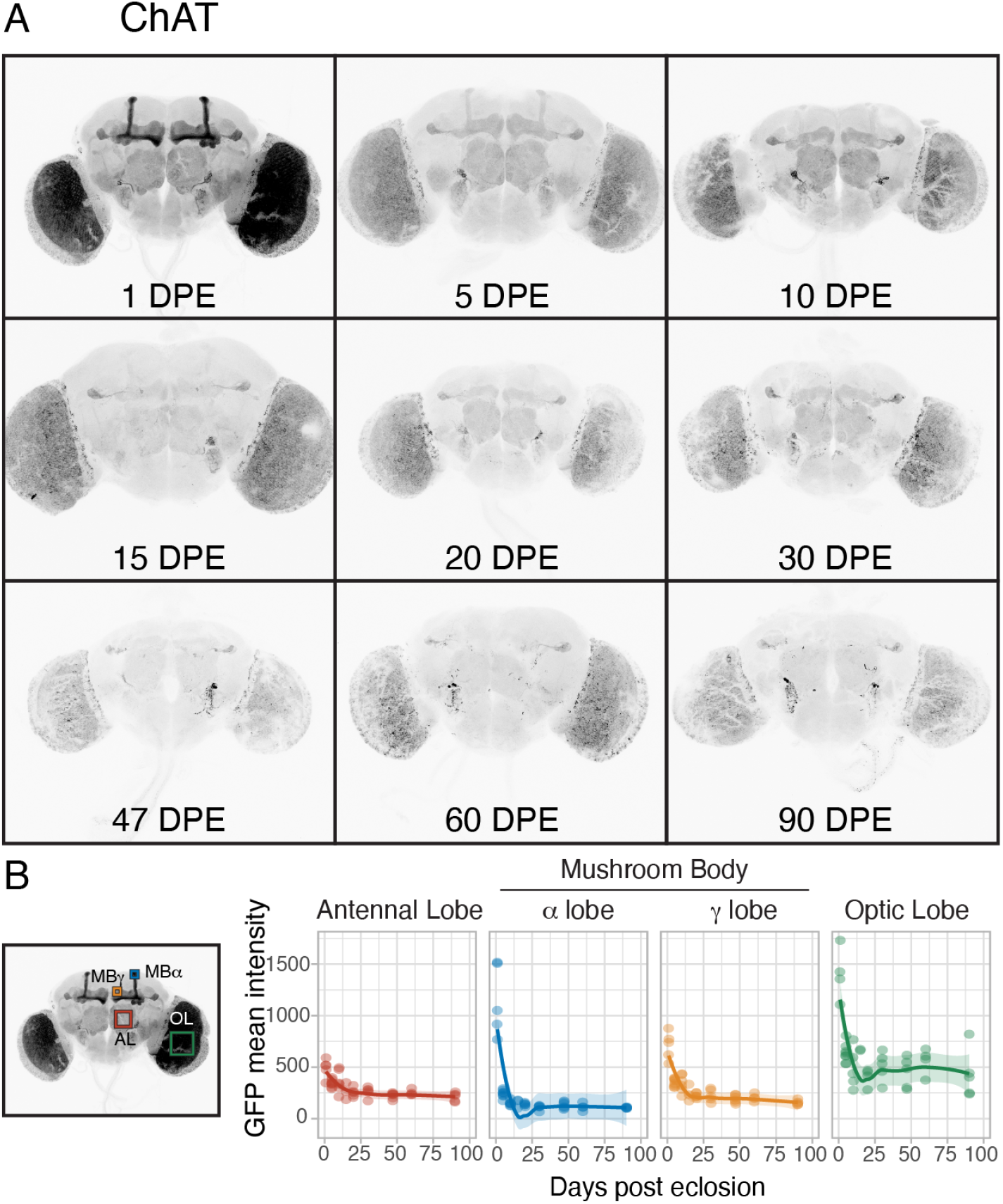
(A) Flies expressing membrane-bound GFP under the control of *ChAT-GAL4* and a temperature-sensitive GAL80 were induced for 24-hour time windows by shifting from 18°C to 29°C at nine time points throughout adult life (up to 90 DPE). (B) Four brain regions were chosen for quantification of GFP mean intensity (n = 4-8; except 15 DPE, with only three brains).

*ChAT-GAL4* offered the most striking change in expression pattern over the few days post-eclosion, most notably in the optic lobe, where GFP intensity decreased to almost a third of the 1-DPE baseline (Fig. 4). The mushroom body transitioned from a strong predominance of alpha/beta lobes (1 DPE) to an equal level of both alpha/beta and gamma lobes (5 DPE), followed by a stage where only the gamma lobes had detectable signal (30 DPE). The rest of cholinergic brain areas seemed to follow a gradual decline in driver expression; the optic lobes remained the main visible areas at 90 DPE, in addition to the subset of neurons seen with *nSyb-GAL4* and another group of neurons projecting to the mushroom body. We also noticed the seemingly random appearance of individual neurons in older brains with all three drivers (for examples, see arrows in figs. 2A and 3A). We observed a similar expression pattern in 50-DPE brains from uninduced flies, implying the GAL4 activity present in older brains is likely due to a lack of repression from GAL80. This was especially noticeable in the optic lobes for *nSyb-GAL4* (Supplementary Fig. 1).

### Reporter levels decline with age regardless of the targeted insertion site

Previous studies have suggested that differences in expression levels or distribution could be affected by the insertion site of the reporter transgene, raising the possibility that the local chromatin environment of the reporter, rather than activity of the driver, might be affecting an age-related decline in activity (Markstein et al., 2008). To test this, we repeated the same time course with identical *UAS-myrGFP* transgenes inserted into two other targeted insertion site loci at attP40 (chromosome 2L) or su(Hw)attP5 (chromosome 2R). As previously published by the Rubin lab, we found baseline differences in levels and distribution of the membrane-bound GFP reporter among insertion sites at 1 DPE (Pfeiffer et al., 2010), with the most variability observed in the neurons of the optic lobe. However, the insertion sites did not seem to affect the rate of decline over our ageing time course. Our data indicate that the age-related decline in reporter activity stems solely from the GAL4 driver line, at least in neurons (Fig. 5, Supplementary Figs. 2 to 7).

**Figure 5.**
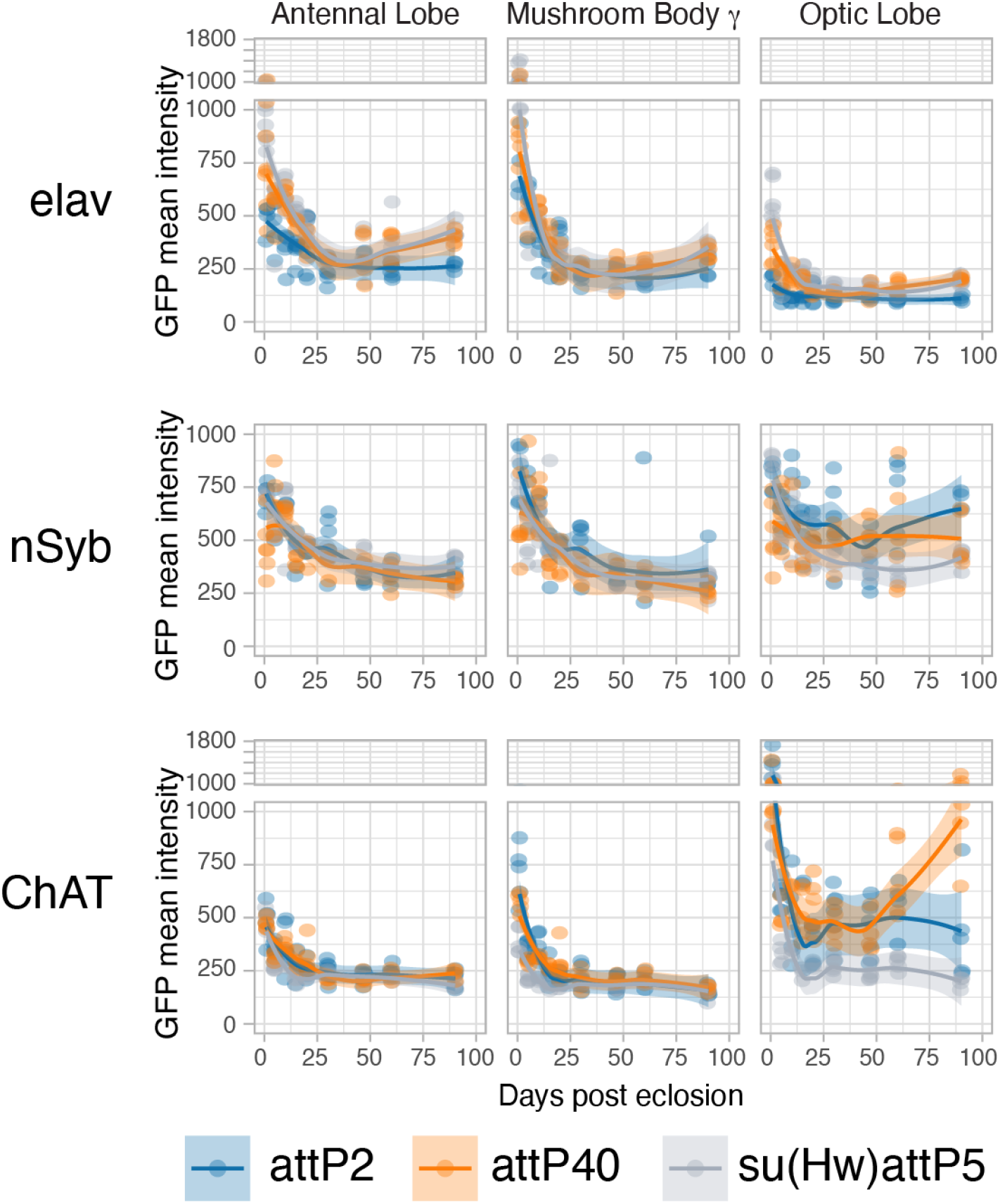
Flies expressing membrane-bound GFP (inserted in attP2, attP40, or su(Hw)attP5) under the control of *elav[C155]-GAL4*, *nSyb[R57C10]-GAL4*, or *ChAT-GAL4* and a temperature-sensitive GAL80 were induced for 24-hour time windows by shifting from 18°C to 29°C at nine time points throughout adult life (up to 90 DPE). Quantification of GFP mean intensity in three brain regions is shown (representative brain images are in Supplementary Figs. 2 to 7).

### Glial driver activity similarly declines during brain ageing

Previous transcriptomic studies have found a decline in expression of neuronal functional genes with ageing, but also a concomitant increase in glial expression of specific genes (for review, see Ham and Lee, 2020). However, a recent study found the activity of two glial GAL4 drivers remained unaltered in older flies (Sheng et al., 2023). To check whether the decline in gene expression is specific to neurons, we used the pan-glial driver *repo-GAL4*, derived from a P-element insertion into the *repo* promoter region, and observed a similar decrease in expression at 10 DPE (Fig. 6). This further supports the need to characterise the spatial and temporal expression pattern of brain GAL4 drivers, regardless of the targeted cell type.

**Figure 6.**
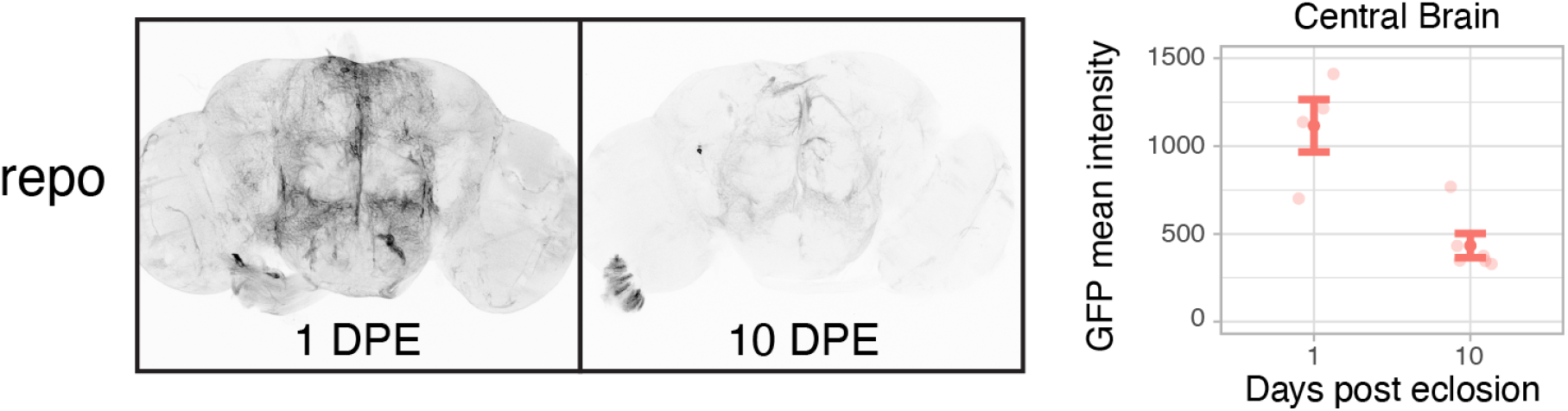
Flies expressing membrane-bound GFP under the control of *repo-GAL4* and a temperature-sensitive GAL80 were induced for 24-hour time windows by shifting from 18°C to 29°C at 1 or 10 DPE. A central brain region was chosen for quantification of GFP mean intensity. Filled circles show average values +/- SEM (n = 4-6).

Taken together, we found that the expression profiles of multiple keystone cell-type defining GAL4 drivers change throughout adult life, both in intensity and region specificity.

## Discussion

GAL4 drivers have provided fly neuroscientists with powerful means to label and control all neurons, with pan-neuronal lines such as *elav[C155]-GAL4* and *nSyb-GAL4*, pan-glial lines such as *repo-GAL4*, or more specific subtypes, such as cholinergic neurons with *ChAT-GAL4*. Even though these drivers have become standards in the field, it is not clear whether they faithfully represent the expected brain expression pattern in ageing flies. We therefore decided to investigate their expression profile over multiple time points in adulthood. We found both pan-neuronal drivers to have distinctly different spatial patterns already in young adults. Moreover, while both showed a bias towards strong mushroom body presence, only *nSyb-GAL4* showed strong expression in most other areas of the brain. The strong differential spatial expression of *elav[C155]-GAL4* came as a surprise since it is the classic driver for pan-neuronal expression (Lin and Goodman, 1994). However, a close look at the literature showed similar images with low signal in the optic lobe compared to the mushroom body (especially the alpha/beta lobes) and antennal lobe (recent examples: Chakravarti Dilley et al., 2020; Hawley et al., 2023; Jonson et al., 2018; Lin et al., 2015). When interpreting published expression patterns of GAL4-driven fluorescent reporters, the use of immunostaining must be taken into consideration as the antibody binding dynamics could alter the signal-to-noise ratio of the reporters and mislead interpretation of subtle differences across brain regions (Wissing et al., 2022). This might contribute to the spatial differences observed in the literature.

In addition to changes in expression patterns during early development, an often-overlooked aspect is how these drivers behave in ageing brains. We found an age-dependent decline in expression activity in all three neuronal and one pan-glial drivers. Characterising these changes becomes crucial as more groups use *Drosophila* to model neurodegenerative diseases. Findings by Seroude and colleagues using a set of 180 GAL4 enhancer trap lines show over 80% of them change expression activity over time as flies age (Seroude, 2002; Seroude et al., 2002). However, it was difficult to find similar studies performed on more widely used neuronal drivers. Only anecdotal findings, such as by the Thor lab, observe a decrease in driving activity of *elav[C155]-GAL4* in older flies (up to 20 DPE; Jonson et al., 2018, 2015); while levels of another pan-neuronal driver (also based on *nSyb*, but derived from a P-element insertion in an unknown locus) remain constant or even increase as flies age. However, these studies did not restrict GAL4 activity to a narrow time window, and because drivers were expressing throughout development, it is difficult to understand the expression dynamics of these drivers. There is also a lack of information in the literature on spatial expression pattern changes in older brains.

The first five days post-eclosion showed the steepest decrease in intensity, with a more gradual decline afterwards. As the temporal restriction requires the flies to be aged at 18°C and therefore to undergo a slower development, we were surprised to observe such changes during the first five days post-eclosion. Many experiments in young adult flies are performed within this time frame; however, most articles report the expression pattern of a specific driver in young flies without temporal restriction. By temporally restricting expression, we were able to reveal spatial pattern changes in the *ChAT-GAL4* and *elav[C155]-GAL4*-driven brains: most notably, a dramatic decrease in expression specifically in the mushroom body (for both drivers) and the optic lobe (for *ChAT-GAL4*). Overall, *nSyb-GAL4* seemed the most consistent driver, with signal still present in most brain regions at 20 to 30 DPE. However, after 30 DPE, even *nSyb-GAL4* expression was significantly reduced. Therefore, we suggest that experiments using GAL4-driven models in older flies need to be interpreted carefully, as most phenotypic changes may be a consequence of expression during development and immediately post-eclosion. Moreover, the spatial changes observed over the first days post-eclosion, for example across mushroom body lobes, could complicate the interpretation of experiments aiming to understand the behavioural role of a specific cell type.

Every driver we have checked so far has shown a decline in expression over time, correlating with previous transcriptomic studies reporting a general age-dependent decrease of gene expression, especially in neurons (Berchtold et al., 2008; Erraji-Benchekroun et al., 2005; Lu et al., 2004). However, measuring the relative levels of transcripts could lead to misinterpretations because it relies on the assumption that overall RNA levels across samples are similar (Lovén et al., 2012). Our results provide an alternative approach to confirm the decrease of absolute gene expression at a region-specific level.

The difficulty to find a single GAL4 line exhibiting a consistent expression level in the ageing brain likely stems from the fact that most cell-type specific drivers by nature must rely on the enhancer regions of genes important for development or function, most likely to be downregulated in older brains. Additionally, a single-cell transcriptomic study found that the expression of genes commonly used to annotate cell types also change throughout ageing (Ximerakis et al., 2019). Thus, it seems unlikely that we could find a single enhancer region to drive both cell specificity and temporal stability during ageing. A possible approach could be an intersectional method where one component provides the cell specificity while the other allows for a consistent expression over time. The crucial step would be finding a gene, most likely part of the cellular housekeeping machinery, that is consistently expressed in a majority of cell types as the brain ages. Promising examples are genes encoding components of the ribosomal machinery, which were found by single-cell RNAseq to be expressed at high levels in older brains and part of a core group of genes expressed in most cell types (Davie et al., 2018; Yang et al., 2022).

An additional point that was not addressed in our study is the effect of ageing on potential ectopic expression changes of these drivers. As we have focused on the brain, we cannot rule out changes in other tissues, which could have additional effects in assays testing behaviours such as survival or locomotor activity. To the best of our knowledge, we have not found reports other than from the Seroude group investigating ectopic expression of GAL4 drivers in ageing (Seroude et al., 2002).

Unexpectedly, we also found aberrant expression of these drivers in uninduced 50-DPE brains, even though there is very little uninduced expression in 1-DPE brains, revealing limitations in the use of temperature-sensitive GAL80 to repress GAL4 activity in older flies. One probable explanation is that the promoter used to drive GAL80[ts], which is derived from the *alphaTub84B* gene, may not be ubiquitous across the brain during ageing. An intriguing recent report described the appearance of GFP-expressing clones in larval brains even in the presence of GAL80, a phenomenon of gene expression plasticity (coined “Illuminati”) that seems restricted to the brain (Goupil et al., 2022). A dysregulation of this process could potentially explain the aberrant expression we observe in older brains. An alternative method to temporally restrict GAL4 activity is GeneSwitch, where expression is turned on by adding RU486 to the food (Osterwalder et al., 2001). However, characterisation of a GeneSwitch cassette downstream of the *elav* promoter region showed an age- and sex-dependent expression variation upon induction, in addition to a leaky background during development, making this system a yet less reliable alternative (Poirier et al., 2008).

To the best of our knowledge, this is the first study investigating the expression profiles of commonly used brain GAL4 drivers over multiple temporally restricted time points in ageing brains. Our findings highlight the variability of their spatio-temporal activity, even in young adult flies. Out of the two pan-neuronal drivers we profiled, *nSyb-GAL4* had the most consistent expression profile across the brain in adult flies, making it a preferable GAL4 for long-term pan-neuronal expression in adults. However, all drivers were very weak after only 30 DPE at 18°C. This lack of reliable driver activity later in life implies phenotypes observed with current neurodegenerative disease models may be mainly due to the effect of a transgene during development or soon after eclosion, and results need to be interpreted accordingly. To study the interaction of ageing and neurodegeneration, where the protein of interest is only produced in older brains, new driver systems will need to be developed. Our results also have implications to the broader fly community, as they underscore the critical importance of characterising GAL4 driver lines in detail during experimental design, and we hope this study will encourage other researchers to add to this resource.

## Acknowledgements

We thank Jake W. Newland for helpful comments on the manuscript. This work was supported through NHMRC grants APP1128784 and APP1185220, Ian Potter Foundation grant 20190091, and philanthropic funding from the Menzies Institute for Medical Research, to OJM.

## Supplementary figures

**Supplementary Figure 1.**
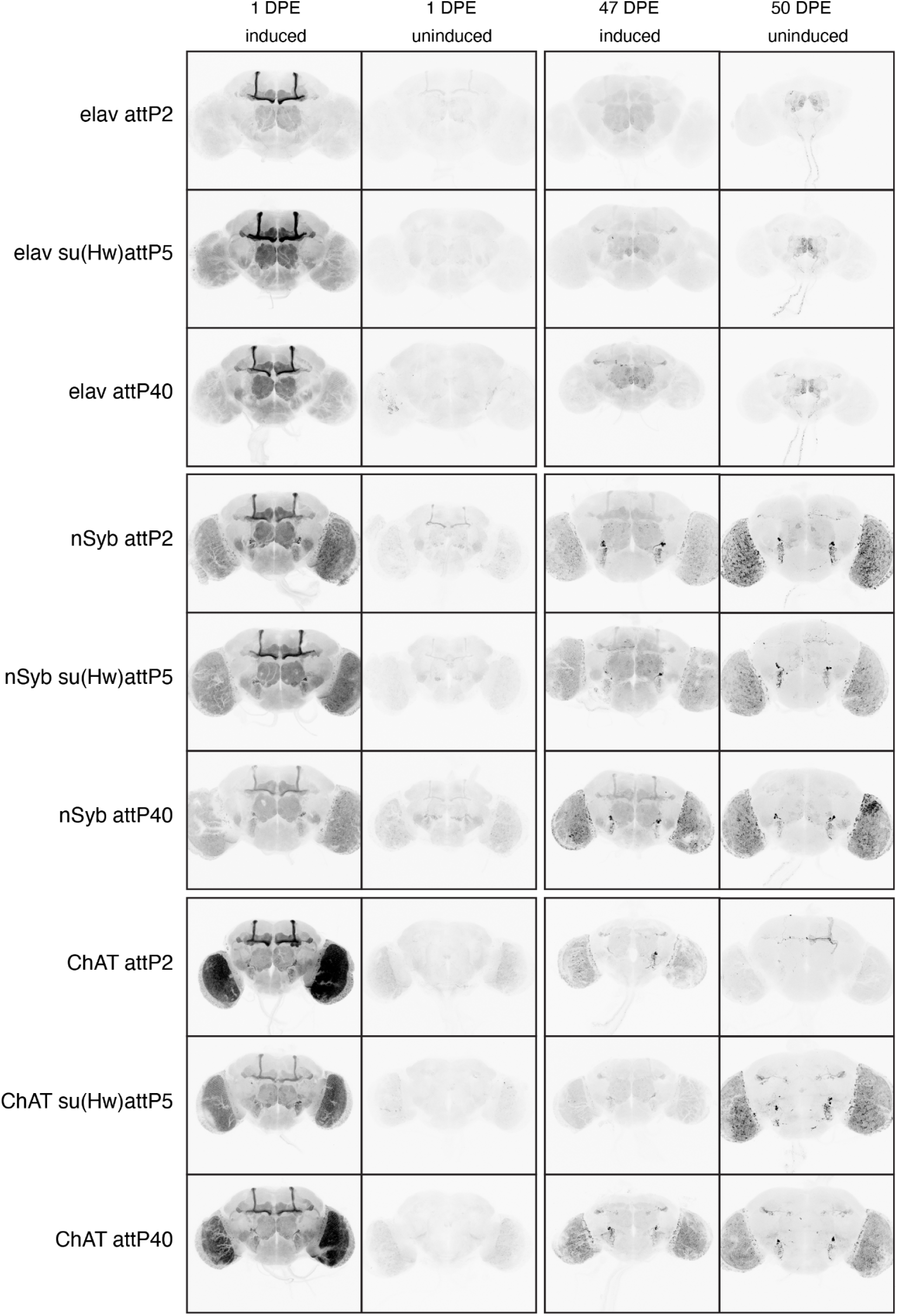
Comparison between brains from flies that were induced at 29°C for 24 hours at 1 or 47 DPE and flies that were left uninduced at 18°C until 1 or 50 DPE.

**Supplementary Figure 2.**
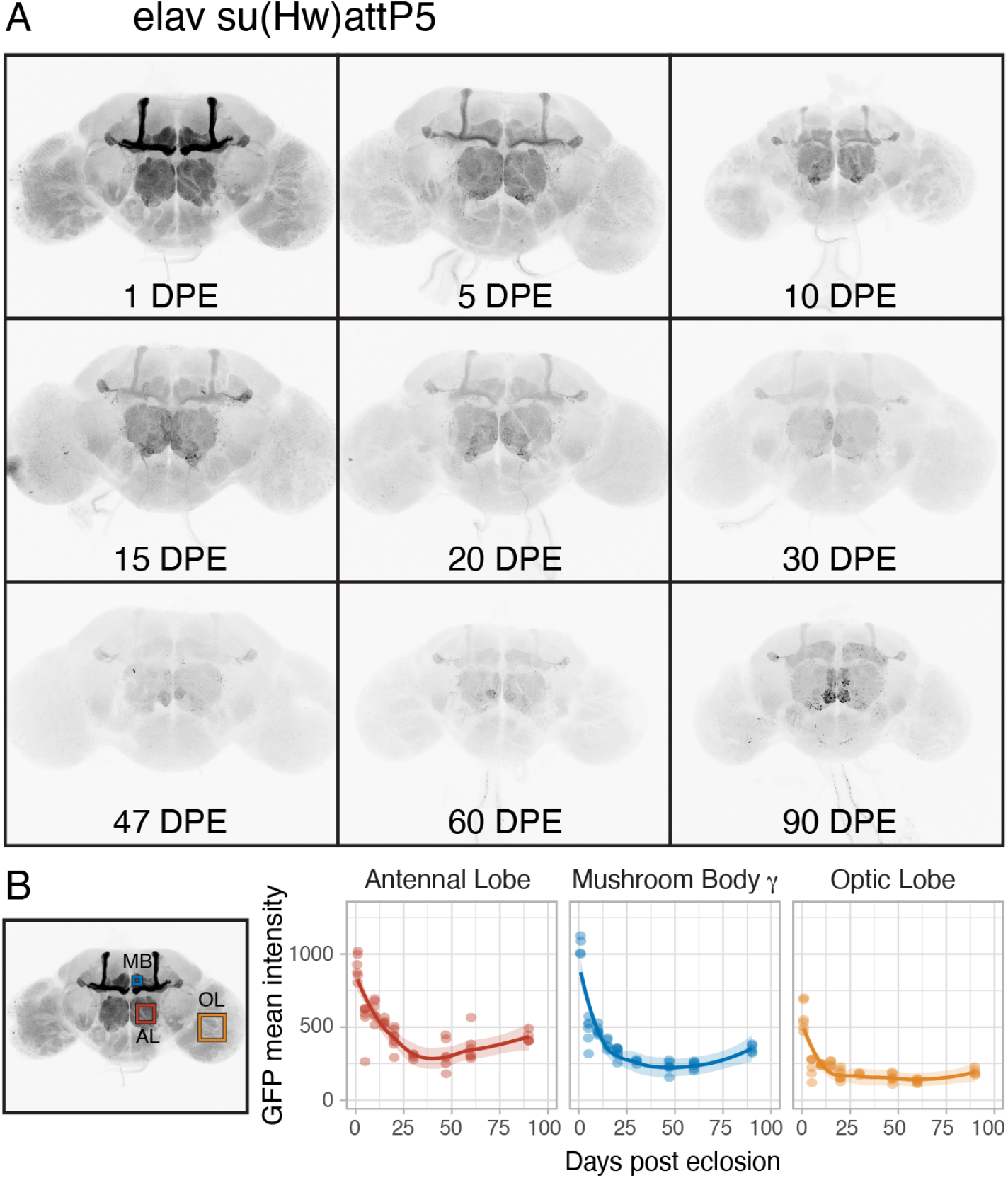
(A) Flies expressing membrane-bound GFP (inserted in su(Hw)attP5) under the control of *elav[C155]-GAL4* and a temperature-sensitive GAL80 were induced for 24-hour time windows by shifting from 18°C to 29°C at nine time points throughout adult life (up to 90 DPE). (B) Three brain regions were chosen for quantification of GFP mean intensity.

**Supplementary Figure 3.**
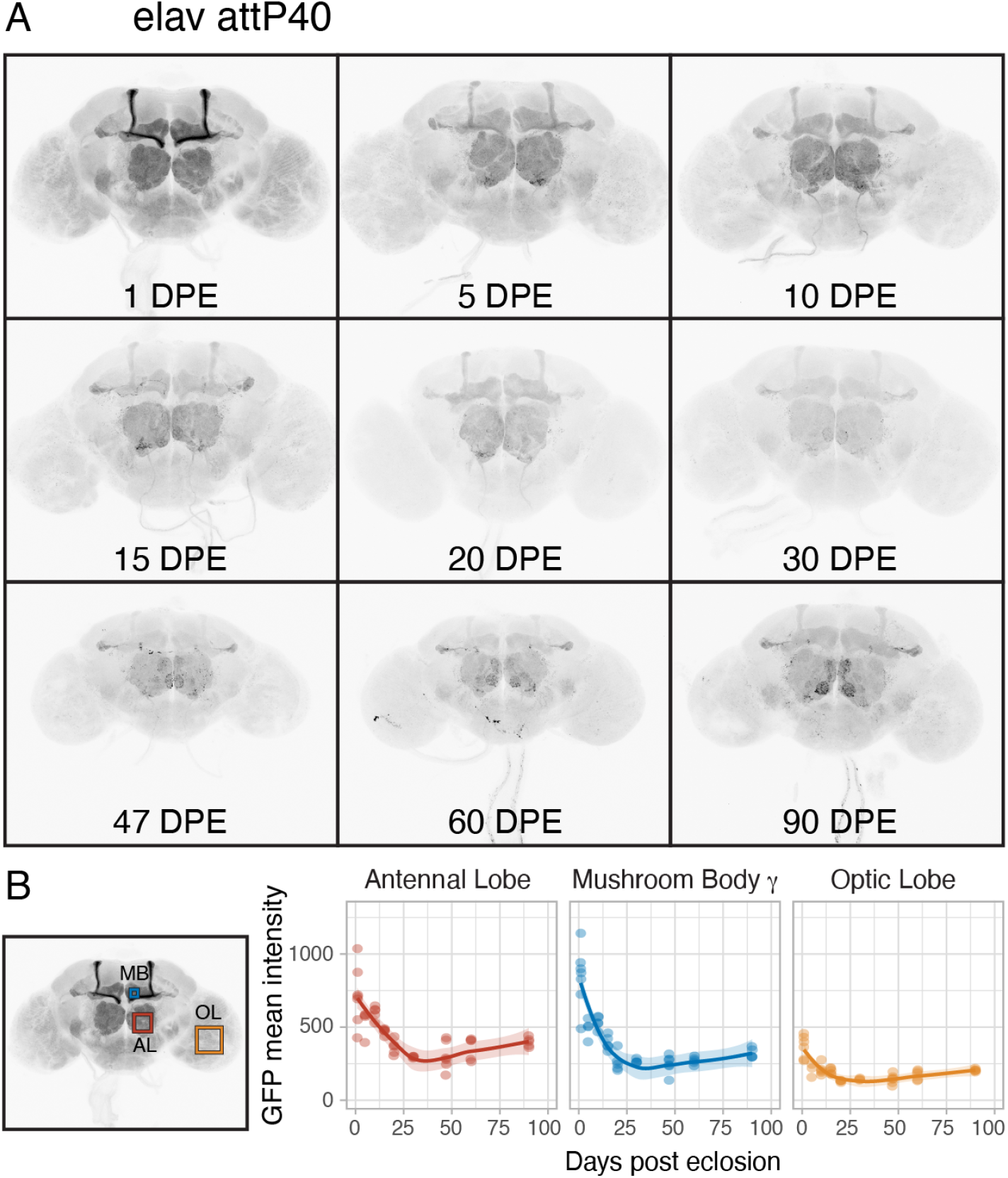
(A) Flies expressing membrane-bound GFP (inserted in attP40) under the control of *elav[C155]-GAL4* and a temperature-sensitive GAL80 were induced for 24-hour time windows by shifting from 18°C to 29°C at nine time points throughout adult life (up to 90 DPE). (B) Three brain regions were chosen for quantification of GFP mean intensity.

**Supplementary Figure 4.**
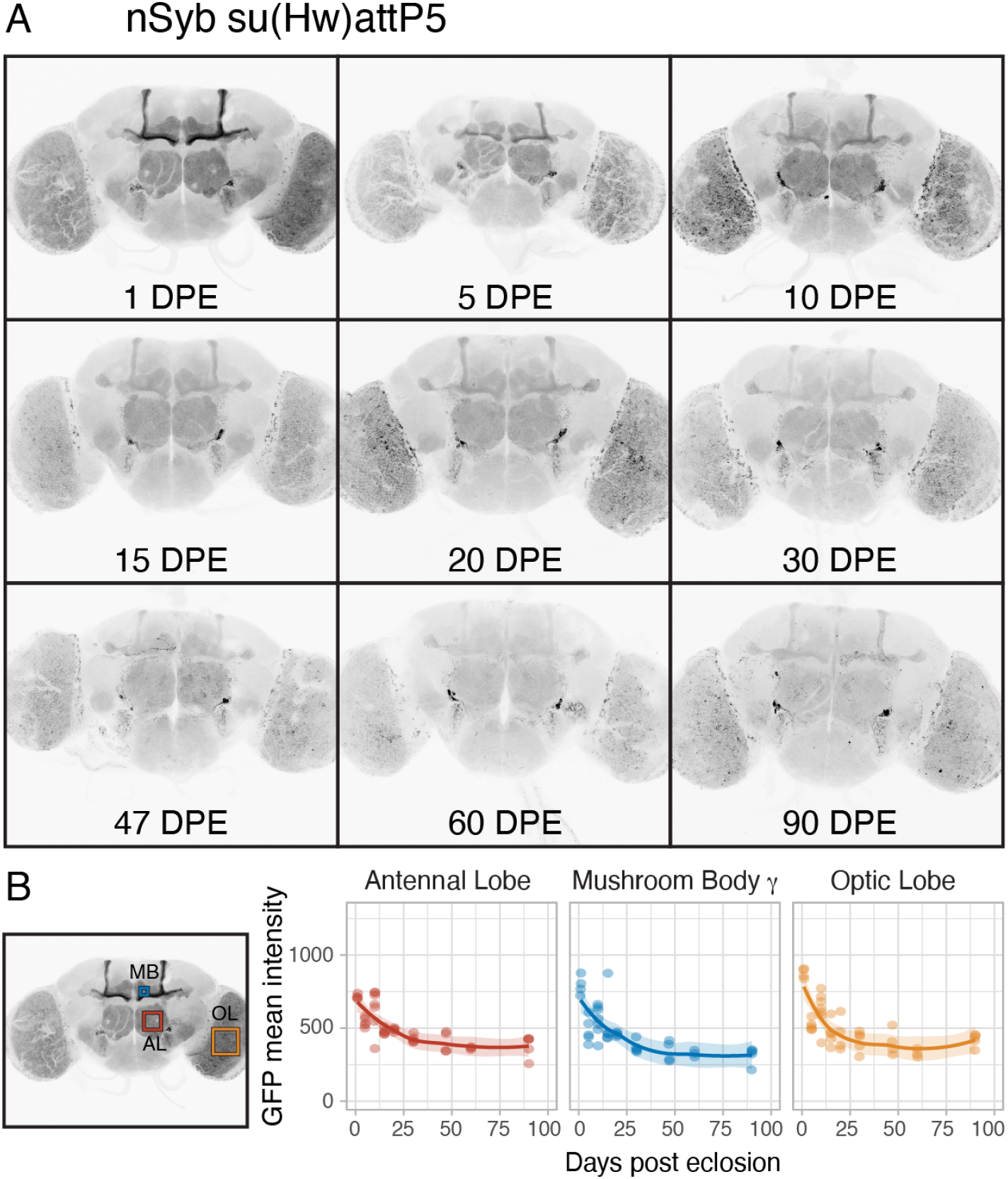
(A) Flies expressing membrane-bound GFP (inserted in su(Hw)attP5) under the control of *nSyb[R57C10]-GAL4* and a temperature-sensitive GAL80 were induced for 24-hour time windows by shifting from 18°C to 29°C at nine time points throughout adult life (up to 90 DPE). (B) Three brain regions were chosen for quantification of GFP mean intensity.

**Supplementary Figure 5.**
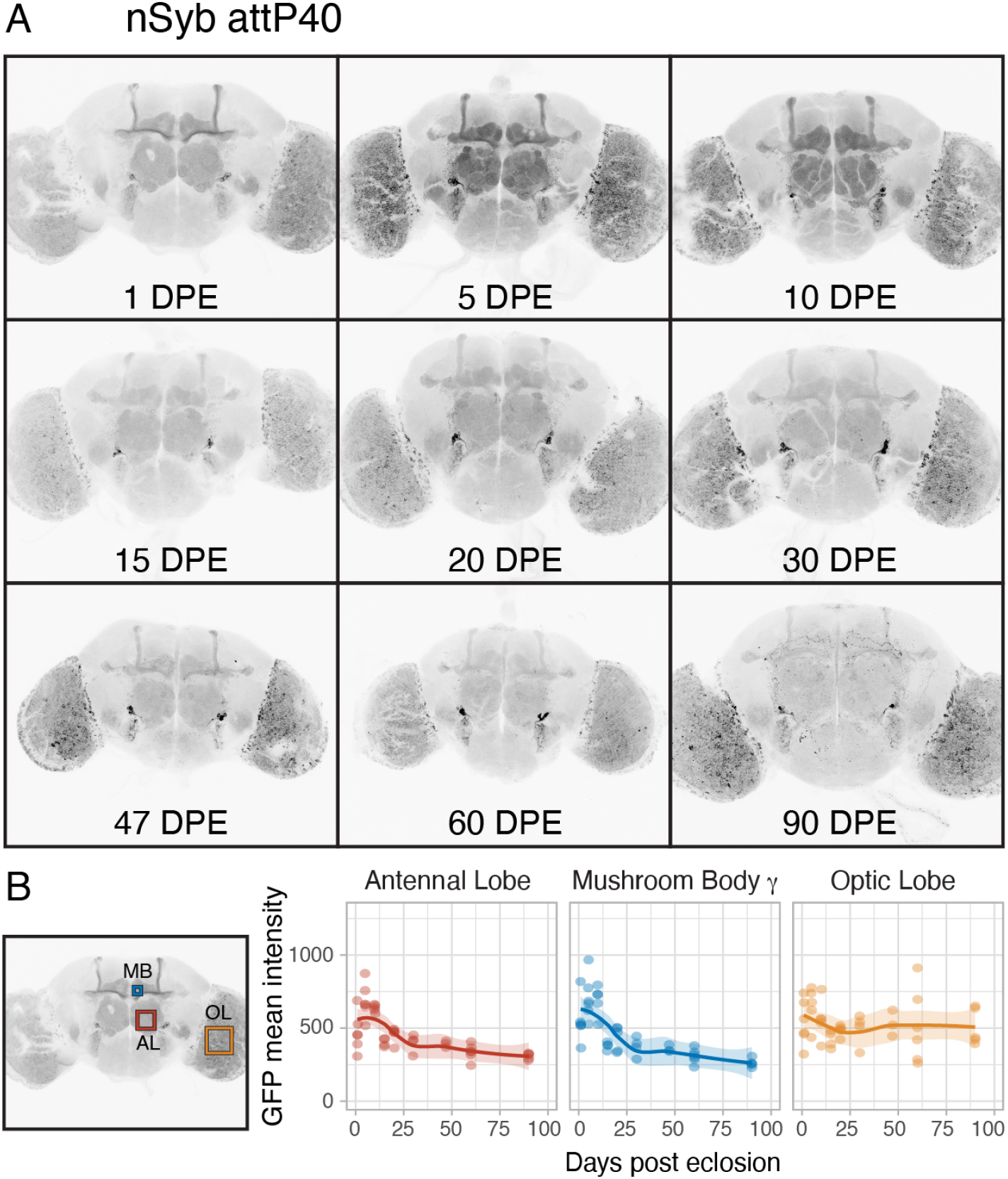
(A) Flies expressing membrane-bound GFP (inserted in attP40) under the control of *nSyb[R57C10]-GAL4* and a temperature-sensitive GAL80 were induced for 24-hour time windows by shifting from 18°C to 29°C at nine time points throughout adult life (up to 90 DPE). (B) Three brain regions were chosen for quantification of GFP mean intensity.

**Supplementary Figure 6.**
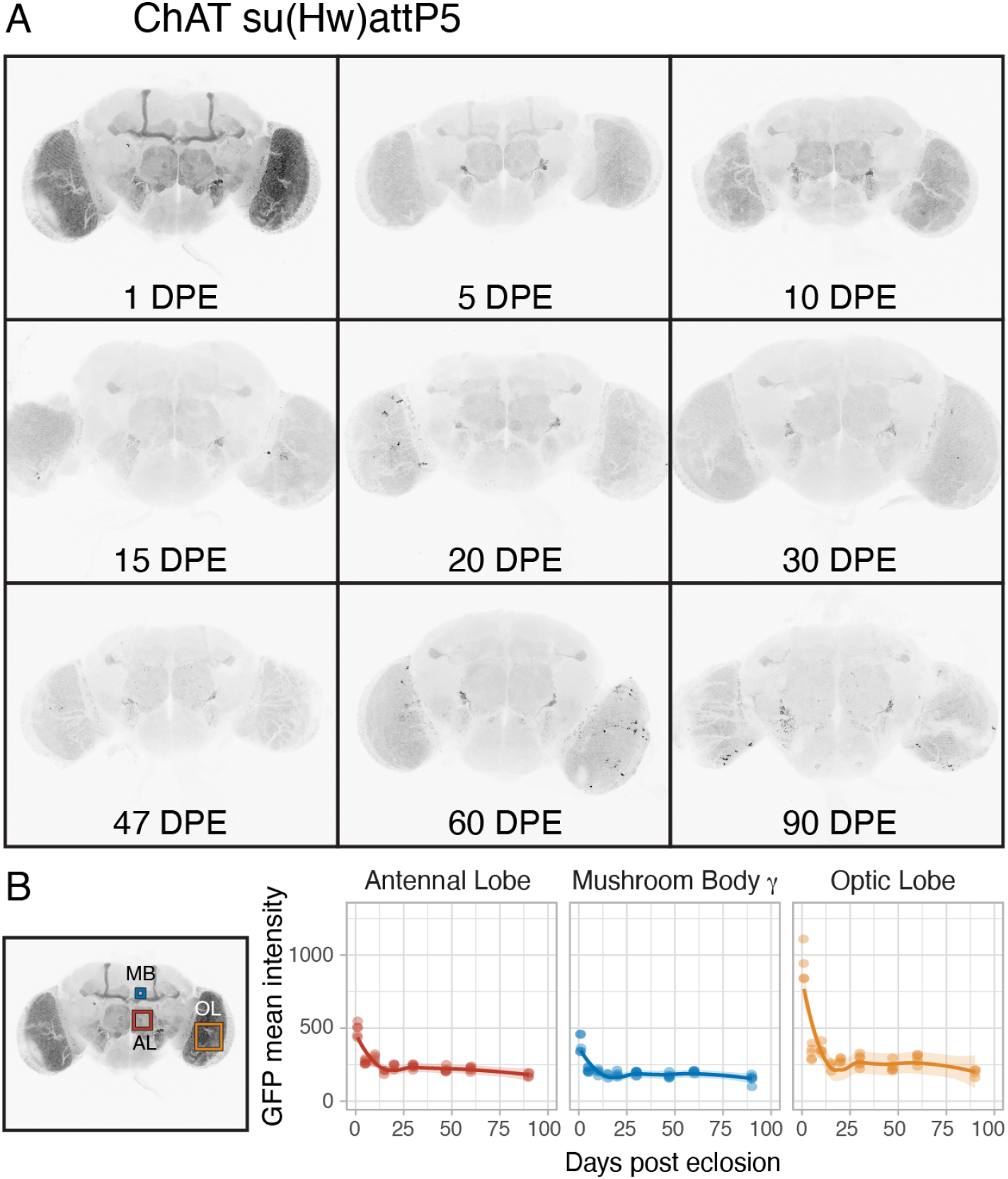
(A) Flies expressing membrane-bound GFP (inserted in su(Hw)attP5) under the control of *ChAT-GAL4* and a temperature-sensitive GAL80 were induced for 24-hour time windows by shifting from 18°C to 29°C at nine time points throughout adult life (up to 90 DPE). (B) Three brain regions were chosen for quantification of GFP mean intensity.

**Supplementary Figure 7.**
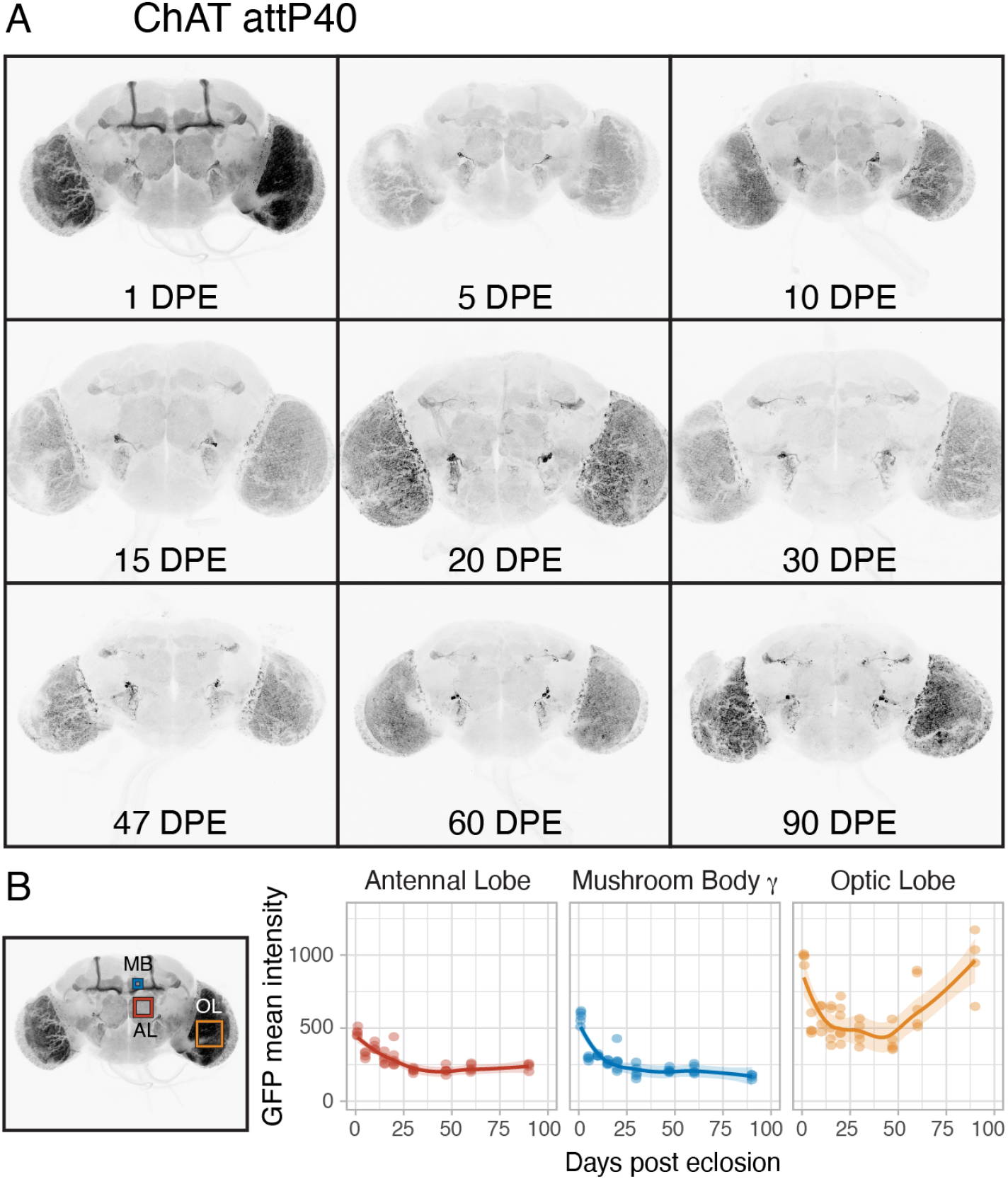
(A) Flies expressing membrane-bound GFP (inserted in attP40) under the control of *ChAT-GAL4* and a temperature-sensitive GAL80 were induced for 24-hour time windows by shifting from 18°C to 29°C at nine time points throughout adult life (up to 90 DPE). (B) Three brain regions were chosen for quantification of GFP mean intensity.

